# qDATA - an R application implementing a practical framework for analyzing quantitative Real-Time PCR data

**DOI:** 10.1101/2023.11.29.569183

**Authors:** Adrian Ionascu, Alexandru Al. Ecovoiu, Mariana Carmen Chifiriuc, Attila Cristian Ratiu

**Affiliations:** Drosophila Laboratory, Faculty of Biology, University of Bucharest, 060101 Bucharest, Romania; Department of Botany and Microbiology, Faculty of Biology, University of Bucharest, 060101, Romania; The Research Institute of the University of Bucharest, Faculty of Biology, University of Bucharest, 050095 Bucharest, Romania

**Keywords:** qRT-PCR, statistics, data analysis, bioinformatics software, R programming

## Abstract

Gene expression assays that are based on quantitative real-time PCR (qRT-PCR) method are still very popular, therefore, we developed qDATA, a modern open-source R based bioinformatics application that offers a quick and intuitive analysis of raw cycle threshold (Ct) values. The application relies on a straightforward specific data input consisting in Ct values and a few other mandatory fields specifying the experimental and control groups, as well as the target and house-keeping genes. Afterwards, it automatically performs descriptive statistics, normality and statistical testing on 2^-ΔCt^ (or ΔCt) and 2^-ΔΔCt^ terms calculated with Livak’s method. Each computation is indicated in dedicated tabs that display their output consisting in dynamic data tables and publishing quality graphs. We also detail a qRT-PCR data analysis framework that depends on performing exhaustive ΔCt calculations within discrete biological replicates (BRs). The Livak formula is used for the complete sets of available data. These prerequisites arguably lead to an improved data analysis and statistical relevance. In order to maximize the relevance of the results we recommend using at least three BRs and three technical replicates. The efficiency of our computing approach was tested with qDATA using input Ct values corresponding to immune related gene expression evaluated in experimental infection of *Drosophila melanogaster* and summer versus winter *Apis mellifera* workers, respectively. Our results reveal that our working strategy is very reliable and highlight the efficacy and performance of qDATA application.

## 1. Introduction

Functional genomics focuses on genes and their regulation, a highly complex process that tunes the timing and the amplitude of gene expression. Nowadays, genome-wide gene expression is assessed by using sophisticated highthroughput methods, such as microarray and RNA sequencing. Regardless of the massive amount of data produced by these technics, they are still deficient in providing accurate measurements of gene expression fold-changes (FC). The quantitative real-time PCR (qRT-PCR) is a very robust method that uses fluorescence in order to monitor the specific DNA accumulation through successive rounds of amplifications. Although somehow challenging because it requests accurate pipetting steps, qRT-PCR delivers more precision than the alternative highthroughput methods, and therefore is still considered the gold-standard for quantifying gene expression [1]. The specific output data for qRT-PCR are the so-called cycle threshold (Ct) values or raw Ct values that are often exploited to calculate FC values (2^-ΔΔCt^ values) by using Livak’s method [2] which assumes equal PCR amplification efficiencies. The Livak formula implies that the Ct values specific for a housekeeping (HK) gene or endogene are subtracted from the Ct values corresponding to a target gene as a form of data normalization. The common practice of qRT-PCR data normalization based on single housekeeping gene was analytically argued elsewhere [3]. As far as we know, a factual routine for effectively using the Ct values in Livak mathematical equations is not openly addressed but instead it is somehow assumed that the qRT-PCR data analysis approach is a common knowledge. Even if guidelines for maximizing the accuracy of qRT-PCR data analysis are periodically updated [4] the Livak approach is still widely used.

As it was not in our intention to debate neither the inherently natural variability of the living tissues considered for genes expression assessment, nor the origin of various technical errors impeding on the results, we focused on the automatization of a straightforward methodology for the analysis of qRT-PCR data. Our data analysis framework considers at least three biological replicates (BRs) and three technical replicates (TRs) per each BR. The associated qRT-PCR data are interpreted by our original bioinformatics software in order to obtain robust conclusions about their statistical and biological significance. Our tool computes and then employs all of the alternative ΔΔCt values (and their 2^-ΔΔCt^ counterpart values) and relies on either the raw ΔCt values or on their linearized form for running specific statistics tests.

Herein, we present the functionality of qDATA, an original open-source R application adorned by a user-friendly interface. qDATA is able to accommodate various qRT-PCR datasets for statistical analysis according to a methodology which is further detailed.

## 2. Implementation

### 2.1. qRT-PCR data analysis framework

The paradigm of the modern experimental sciences is the reproducibility of the data obtained in similar or nearly identical conditions. Accordingly, qRT-PCR studies are usually performed using at least three BRs (samples) and each of them is inquired by three TRs. This experimental approach is conceived to identify inherent procedural and/or technical errors. Consequently, if one gene of interest (GOI) and one endogene are considered, for each BR six Ct values are to be generated, namely three for the GOI and other three for the endogene (Figure 1).

**Figure 1.**
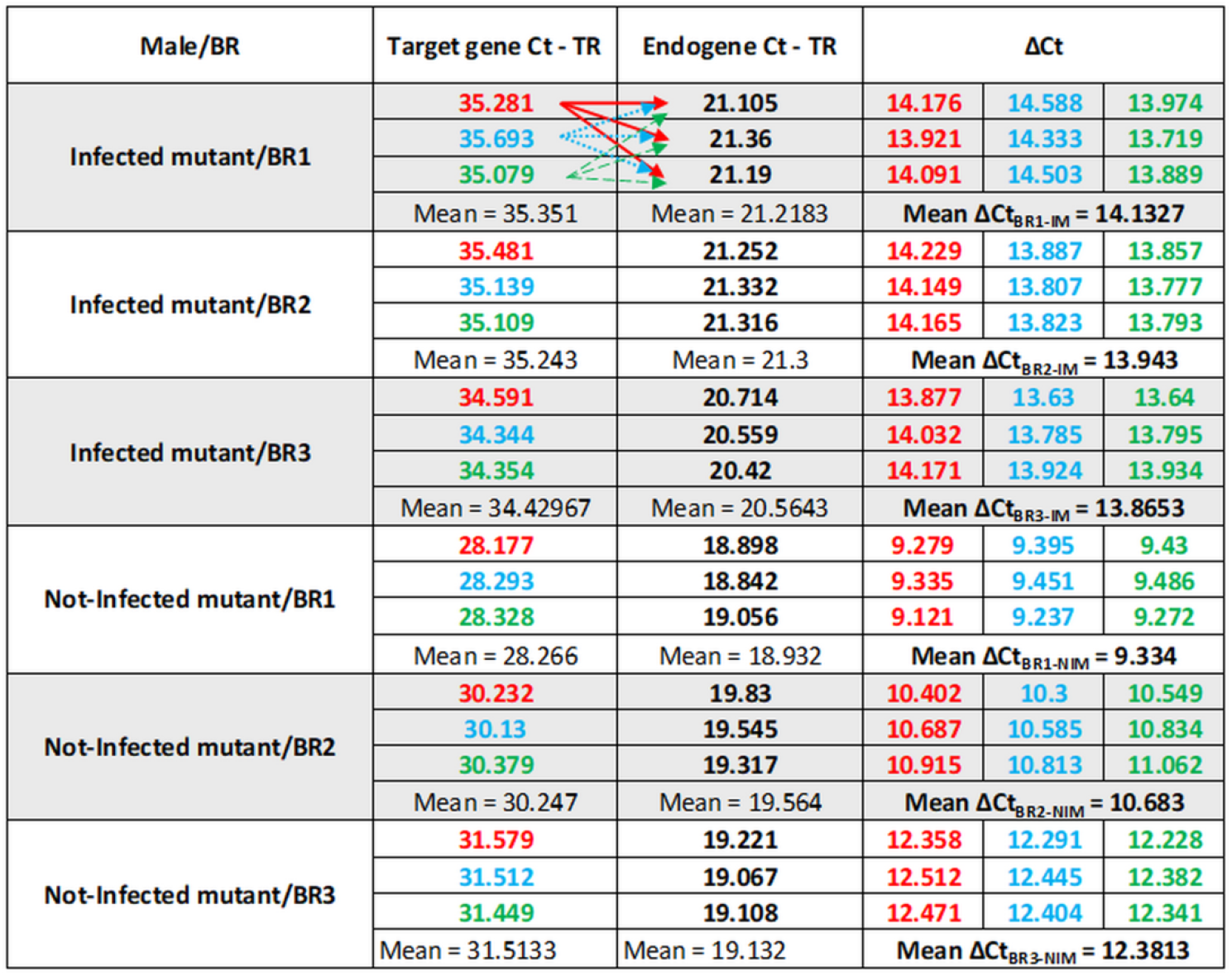
Tabular data showing a standardized procedure for interpreting triplicate Ct data (TRs) collected for a qRT-PCR performed on *LManVI* target gene. Three distinct BRs from either infected or not-infected *γCOP*^*14a*^/ *γCOP*^*14a*^mutants were inquired. According to Livak’s formula, ΔCt values are obtained by subtracting the endogene Ct values from the target Ct values. Considering a single BR, a total of nine ΔCt values result by subtracting each endogene Ct from each of the target Ct. The arrows (shown only for the Infected mutant/BR1 category) point to the values that are to be subtracted from each target gene Ct. Alternatively, a mean ΔCt can be calculated for each BR, either by averaging the nine ΔCt values or by subtracting the mean endogene Ct from the mean target Ct. By red, blue and green numbers we indicated the groups of ΔCt values originating from a given target Ct. IM and NIM acronyms indicate the infected mutant and, respectively, not-infected mutant BR.

In figure 1, we present Ct data obtained during a series of studies that were previously published by our laboratory. Briefly, groups of *Drosophila melanogaster* males homozygous for a mutant allele of *γCOP* gene, symbolized *γCOP*^*14a*^ [5] were infected with a *Pseudomonas aeruginosa* bacterial inoculum delivered by ingestion [6]. A non-infected control group of *γCOP*^*14a*^/*γCOP*^*14a*^ males was also reared in the same experimental conditions. For the assessment of the expression changes of *Lysosomal α-mannosidase VI* (*LManVI*) target gene consecutive to the experimental bacterial infection, the respective Ct data were normalized relative to *Ribosomal protein L32* (*RpL32*) endogene [7].

Thus, for each BR or sample, three distinct Ct measurements were performed for the target gene and, respectively, the endogene. For generating ΔCt values (data normalization), there are two alternative approaches. One is to subtract average Ct values corresponding to endogene from the average Ct values of target gene, which leads in the case of Infected mutant/BR1 (IM/BR1) to a mean ΔCt = 14.1327 (Figure 1). This approach is rather frequently used and its line of reasoning is that there is no apparent reason to pair a specific target Ct value with any particular endogene Ct value, since each reaction takes place in a distinct well. Such an approach is possible when real time PCR methods based on SYBR Green chemistry are considered. If both the target and endogene Ct are collected by reading a single well, the pairing of those values is implicit. Nevertheless, this logic does not exclude a strategy where every possible ΔCt values are calculated by coupling every single target Ct to each endogene Ct from within the same sample.

Here, we argue that any calculations of the ΔCt values should be performed only with data gathered for a single BR, resulting in nine ΔCt values. Although very often the BRs are highly comparable, they have inherent specificities, especially at molecular level, that require to be distinctly addressed. This strategy provides nine ΔCt values with a mean of 14.1327, as expected, but also allows estimating the data intrinsic variability, which often is measured as the differences between each value and the sample mean. In other words, the distribution of a data set contains a certain degree of uncertainty, frequently expressed as mean ± standard deviation (SD), which is further used for the standard error (SE) computations. Our view is that the estimated variance of a data set should be carried through all the calculations of Livak’s method [2]. For example, if the error estimates are to be considered, the mean target Ct value from IM/BR1 would be 35.351 ± 0.3129, while the average endogene Ct would be 21.2183 ± 0.1298. When the rules for propagation of errors are applied, the corresponding ΔCt value is 14.1327 ± 0.3387. Alternatively, the uncertainty estimated for the mean of the nine possible ΔCt values within the same sample gives 14.1327 ± 0.2934. In this case, the small difference between the two SD values probably is not critical, but pinpoints two possible limitations: (i) the error estimates will have other values when using data set means than when using the full set of raw Ct for inferring the mean; (ii) straight forward calculation sheets, such as those offered by Microsoft Excel or LibreOffice Calc, do not implicitly consider the error propagation, therefore, in some specific situations, the regular user could misevaluate the final results.

The ΔΔCt critical value is influenced by the strategy chosen to generate it, as for example if using the mean ΔCt value or all of the individual ΔCt values. Moreover, handling of the ΔCt values is of key importance for statistical testing of the differences between the analyzed sample groups. If we consider the *LManVI* gene expression variation among the IM and not-infected mutant (NIM) groups, a specific mean FC value is obtained by applying the Livak method. Regardless of how many individual FC values are computed, the respective data set can be merely submitted to descriptive statistics. Only the ΔCt data sets can be used to evaluate the statistical significance of the differential expression of *LManVI* in IM and NIM individuals. The results provided by the specific statistical tests are dependent on the sample variation; therefore, one has to work with the best available data sets. In our example, by using mean ΔCt values we are able to define discrete data sets with three entries for IM and NIM (because there are three BRs, each with one mean ΔCt). In contrast, if all the 27 possible ΔCt values are computed, the corresponding data sets are more amenable to statistical analysis. We could not find an accepted consensus regarding the use of the ΔCt values for statistical purposes, but it is questionable if they should be used per se. In their seminal paper, Livak and Schmittgen (2001) argue that employing the raw Ct values miss the real variation within a given data set [2]. Instead, their conversion to the linear form using the term 2^-Ct^ leads to a more accurate depiction of data set variation. We consider that dealing with the sets of ΔCt values is an analogous situation, therefore we recommend to use for statistic testing the linear form of ΔCt via the term 2^-ΔCt^. Arguably, this approach grants a bigger consistency for the results of data analysis.

Using our example data, testing the statistical significance of FC differences on the complete set of 27 possible 2^-ΔCt^ versus using only three element sets minimizes the impact of possible outlier Ct values and results in a more refined statistical relevance (Table 1).

**Table 1.** Sample statistics and statistical hypothesis testing performed on different size data sets comprising of 2^-ΔCt^ values specific for *LManVI* gene expression in I and NI mutant males.

| Sample statistics | IM | NIM |
| --- | --- | --- |
| Mean $2^{-\Delta Ct}$ (27) | $6.27 \times 10^{-5}$ | $7.86 \times 10^{-4}$ |
| Mean $2^{-\Delta Ct}$ (3) | $6.21 \times 10^{-5}$ | $7.82 \times 10^{-4}$ |
| SD (27) | $9.91 \times 10^{-5}$ | $5.89 \times 10^{-4}$ |
| SD (3) | $5.80 \times 10^{-5}$ | $6.97 \times 10^{-4}$ |
| SE (27) | $1.91 \times 10^{-6}$ | $1.13 \times 10^{-4}$ |
| SE (3) | $3.35 \times 10^{-6}$ | $4.03 \times 10^{-4}$ |
| 95% CI (27) | $[5.88 \times 10^{-5}, 6.67 \times 10^{-5}]$ | $[5.53 \times 10^{-4}, 10.19 \times 10^{-4}]$ |
| 95% CI (3) | $[4.76 \times 10^{-5}, 7.65 \times 10^{-5}]$ | $[-9.51 \times 10^{-4}, 25.14 \times 10^{-4}]$ |
| Unpaired t-test (p value) |  |  |
| 27 vs 27 | < 0.0001 |  |
| 3 vs 3 | = 0.1484 |  |
| Mann-Whitney test (p value) |  |  |
| 27 vs 27 | < 0.0001 |  |
| 3 vs 3 | = 0.1 |  |

When analyzing the data presented in Table 1 one may notice that the mean 2^-ΔCt^ values are very similar, regardless of the considered data sets. Conversely, the other statistics favor the larger 2^-ΔCt^ datasets comprising of 27 values, excepting for the SD (3) value for IM. Since SE values are affected by using only the mean values, this also impacts the 95% confidence intervals (CI), which are sensitive to sample size and data variability. Here, for both IM and NIM categories the 95% CI are noticeable narrower when the larger 2^-ΔCt^ datasets were considered. Data presented in Figure 1 disclose that the Ct values collected for the three BRs of NIM lead to sizable differences between ΔCt sets or their means. These are the very circumstances when the full spectrum of achievable ΔCt data should be exploited. Descriptive statistics also exert a strong influence over the results of the parametric t-test or the nonparametric Mann-Whitney test. Our example of data analysis revealed that only the comparison among the larger 2^-ΔCt^datasets provides statistical significance, irrespective of running parametric or nonparametric tests.

In our example, the gene expression of *LManVI* is downregulated in IM compared to NIM. The actual FC value is conditioned by the way ΔCt terms are used in Livak formula in order to compute the subsequent ΔΔCt values. A straightforward approach is to calculate for the IM or NIM groups a single mean ΔCt value starting from the mean ΔCt values of each BR and to ignore the error propagation. In this case, the mean ΔCt values of 13.9803 and 10.7994 calculated for IM and NIM groups, respectively, conduct to a FC value of 2^-ΔΔCt^ = 2^-3.1809^ = 0.110. The alternative computing that uses two sets of three mean ΔCt values will generate nine ΔΔCt values as there is no preexisting pairing between IM and NIM groups. The mean FC changes to 0.158, with a SE of 0.045 and a 95% CI = [0.113, 0.202]. Finally, if all the 27 possible ΔCt values per global mutant and control groups are considered, the mean FC (calculated from 729 values) is 0.160, with SE = 0.005 and 95% CI = [0.155, 0.165]. Comparing the three mean FCs, it is obvious that the more complex approaches lead to similar values, but care should be taken because the SEs and CIs are different between the two groups. Since smaller SEs and tight CIs imply a higher degree of trust for the mean FC value, we support the use of the latter strategy for computing FC values as it improves the reliability of the reported mean FC values.

### 2.2. qDATA software design

Considering the proposed framework for the analysis of qRT-PCR data, we have implemented a statistical analysis pipeline based upon an original R script and adorned with a modern graphical user interface (GUI) created with the shiny R package. qDATA is freely available on GitHub at https://github.com/A-Ionascu/qDATA Prior to data analysis we highly recommend installing the latest versions of R (currently at version 4.3.2) and RStudio (currently at version 2023.09.1-494). Both R and RStudio have small footprints on the local machines, with minimum system requirements of 1 CPU core and 1 Gb of RAM. Disk memory requirements depend on the installed packages within R, but we recommend at least 1 Gb of free memory. Our bioinformatics application was designed to accommodate a diversity of operating systems (OS), including Windows, MacOS and Debian based Linux distributions, assuming that the R version 4.1.0 is compatible.

The shiny application is suitable for everyday use due to its automatic package requirements retrieval, quick response time and ability to export data reports based on user inputs. In order to run the application, simply open the qDATA.R file and press the “Run App” button. Upon the initial run of the application, all the required packages are downloaded in RStudio, an operation which can last several minutes.

When the application loads, the user is greeted with a modern GUI structured on two windows: “Livak 2^-ΔΔCt^ method” and “About”. Whilst the former is the main data analysis environment, the latter offers information about all the abbreviations used in the application. The main window contains an always-on left side panel and a central area for results output. The left side panel contains a browse button which is used to select the input table from any directory on the local machine. In order to change the input data table, the user simply uploads another table using the “Browse” button and the application retrieves the output for the newly uploaded data.

The qDATA software can be run in the browser by clicking the “Open in Browser” button at the top of the RStudio IDE window. Therefore, the user can use multiple independent instances of the application in its browser mode. If the application is run from the RStudio IDE, the local machine works as the server both for the RStudio window and the browser instances. Closing the RStudio environment automatically disconnects the server and freezes the browser instances. In order to close the RStudio qDATA window, we recommend using the “Stop” button located at the top of the RStudio IDE Console rather than simply closing the running GUI.

qDATA is designed to analyze qRT-PCR data when provided with raw Ct values from gene expression experiments. As previously stated, we recommend using three BRs with three TRs each for every gene of interest. Nevertheless, qDATA can adapt to any number of genes of interest paired with any number of BRs or TRs. Currently, our implementation is designed for gene expression experiments that rely on a single HK gene. The user must create a .csv file containing the raw data similar to the input_model.csv example provided in the archive available on the GitHub page. The input table should contain seven columns of essential experimental information. The “Sample Group” column contains the names of the experimental or control groups, the “Case” and “Type” columns contain the experimental/control tag and, respectively, the GOI/HK tag. The user can define any number of experimental or control tags, their main purpose being to guide the order in which qDATA considers the groups within a comparison. The “Gene” column is to be filled with the corresponding gene names or symbols. Both the “Sample Group” and the “Gene” columns are user-orientated and have minimal impact on data analysis. For the first four columns, consistency is required as the script is case and blank space sensitive. The “BR” and “TR” columns are completed with information regarding biological and, respectively, technical replicates. Within the “Ct” column are raw Ct data that can be imported from the PCR output table and there is no limit for the number of decimals.

Throughout the data analysis process, the user is able to modify parameters from the left side panel with impact on the entire analysis. The available parameters are: selecting the groups of interest using the checkboxes, choosing the type of data to analyze (either linear form of ΔCt values or actual ΔCt values), changing the statistical significance threshold (α), changing the numbers of decimals shown in output tables (not applicable to p values) and choosing the X-axis label orientation for all graphs (horizontal, 30° angle, 45° angle, 60° angle or vertical). When an “Update results” button is present under a parameter, the modification is applied after clicking the button. Adjusting the number of decimals only influences the display but not the calculations of dependent values.

The central area of the “Livak 2^-ΔΔCt^ method” window is divided into eight output tabs, which are briefly described here. From the “Original file”, the user can save the input_model.csv in a chosen directory. Upon loading the input table, a mirror is created in this tab to enhance data visualization. If the software crashes or no output is displayed in this tab, we recommend checking the format of the input table. The exemplified graphical representations correspond to specific qRT-PCR data collected for *Relish* and *Toll* gene expression in summer and, respectively, winter *Apis mellifera* workers [8].

The “Summary statistics” tab shows summary statistics of the calculated 2^-ΔCt^ (default) or ΔCt values corresponding to each group of interest, according to the input selected in the drop-down list. The summary consists of a table with various parameters (mean, median, standard deviation, variance, standard error, 95% confidence intervals, interquartile range, minimum and maximum values), a boxplot (Figure 2A), a violin plot (Figure 2B) and histograms (Figure 2C), which show the shape of the distribution, as well as the localization of mean and median values.

**Figure 2.**
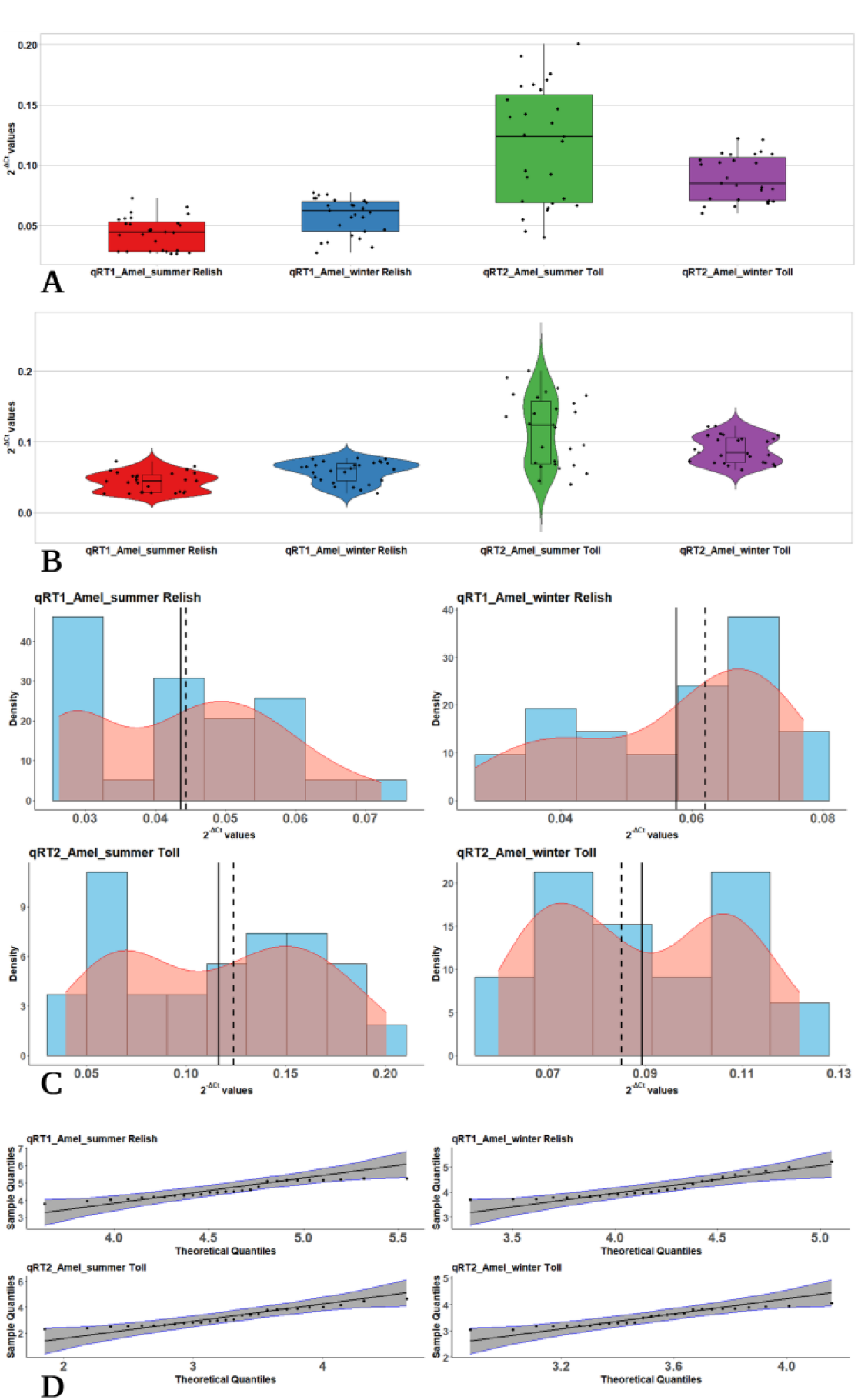
Summary or descriptive statistics performed on 2^-ΔCt^ values derived from qRT-PCR raw data corresponding to *Relish* (qRT1) and *Toll* (qRT2) gene expressions in summer (Amel_summer) and winter (Amel_winter) bees. Information concerning median and mean values, data distribution and shape, as well as its correlation with a normal distribution, are presented using a boxplot (**A**), a violin plot (**B**), histograms (**C**) and quantile-quantile plots (**D**). Individual plots were generated in qDATA and figure edits were created with BioRender.com (accessed on 11 November 2023).

The “Normality statistics” tab displays the results of three tests for data normality (i.e., Kolmogorov-Smirnov, Shapiro-Wilk, Anderson-Darling) performed on the 2^-ΔCt^(default) or ΔCt values. Moreover, quantile-quantile plots (Figure 2D) compare the correlation between sample quantiles and theoretical quantiles of a normal (Gaussian) distribution.

Statistical testing between the 2^-ΔCt^ or ΔCt values corresponding to each group of interest is performed in the “Two sample testing” and “Multiple sample testing” tabs. If intended, both tabs can be customized according to the results displayed on the “Summary statistics” and “Normality statistics” in order to accommodate the most suitable statistical analysis. Three statistical tests with different data normality and equal variances assumptions are available to select. For the “Two samples testing” tab we implemented Student’s two-sample t-test (default), Student’s two-sample t-test with Welch correction and Mann-Whitney U test. For the “Multiple samples testing” tab we implemented one-way ANOVA (default), Welch’s ANOVA and Kruskal-Wallis ANOVA, each paired with a corresponding post-hoc test: Tukey HSD test (default), Games-Howell test and Dunn test respectively. Similarly to post-hoc tests, statistical testing between two experimental groups considers all possible combinations, regardless of biological relevance.

The “Fold Change” tab displays a bar plot and a table based on calculating a dataset of FC (2^-ΔΔCt^) values for comparison between two groups of interest using the Livak method. The plot shows FC values between any given two group with errors bars consisting of one standard error from the mean (Figure 3A). The dotted line marks the threshold of FC = 1, whilst the colored bars indicate upregulation (blue) or downregulation (red). The table contains summary statistics of the FC dataset for each comparison (Figure 3A).

**Figure 3.**
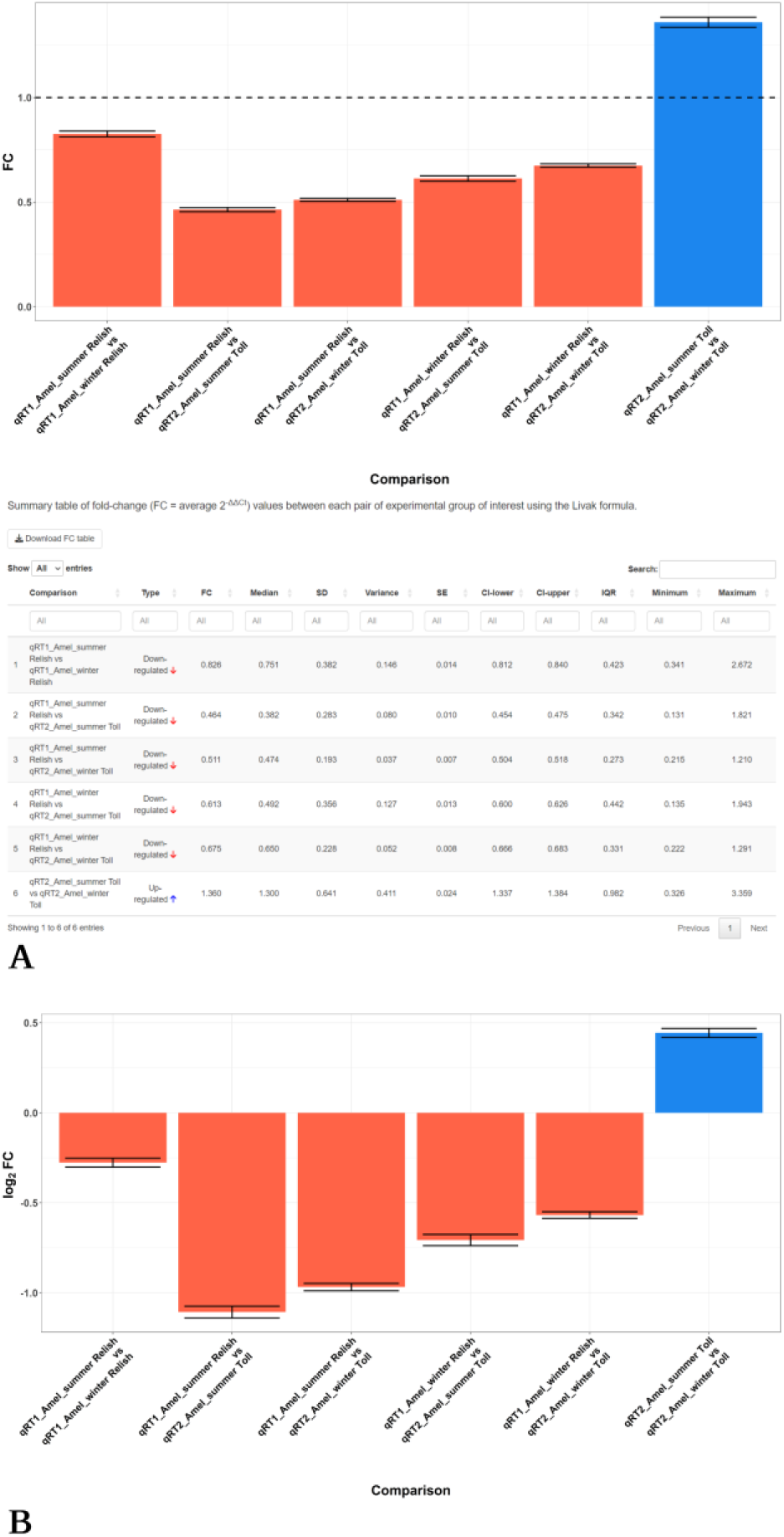
The changes of relative gene expression when every possible comparison is made. The analysis is performed on FC (**A**) and log_2_FC (**B**) values derived from qRT-PCR raw data corresponding to *Relish* (qRT1) and *Toll* (qRT2) gene expressions in summer (Amel_summer) and winter (Amel_winter) bees. The threshold of FC = 1 is indicated by a dotted line; the blue and red bars indicate upregulation or downregulation, respectively. The table showing summary statistics of the 2^-ΔΔCt^ dataset for each comparison is also presented (**A**). Individual plots were generated in qDATA and figure edits were created with BioRender.com (accessed on 11 November 2023).

The “Log_2_Fold Change” tab aims to offer a more intuitive graphical representation of the relative expression due to log_2_ transforming of the mean FC value (Figure 3B). Following this transformation, upregulation will have a positive (blue) and downregulation will have a negative sign (red).

The “Export files” tab contains three additional files, namely the background calculated tables of ΔCt values, the 2^-ΔCt^ values and the FC values. The user can export these tables and use them in other bioinformatics and/or statistical programs.

All displayed tables/plots can be exported by using the button placed above or below the table/plot. The output tables/plots generated by the software consider all of the groups of interest ticked in the left side panel. The name of any exported file will contain a minimal description, including date and time of export, following the format: *table_name/type_day_month_year_hour_minute_second*. Tables are exported in the .csv format and plots are exported in the .png format. The exported tables do not include the GUI implementations, namely the green/red circles standing for the statistical significance and the blue/red arrows for the type of gene regulation.

## 3. Results and Discussion

To evaluate the impact of different possible calculations of ΔCt values on data analysis, we used qDATA for evaluating two types of scenarios or case studies derived for each of the two qRT-PCR raw data sets originally reported elsewhere [6,8]. Case 1 takes advantage of all the available raw Ct values, allowing qDATA to collect all the possible ΔCt values obtained when each target gene TR is coupled with every endogene TR inside any given BR. Therefore, 9 × 3 = 27 ΔCt values are generated for each of the researched categories. We consider that grouped ΔCt calculations within a given BR is a more relevant method of analyzing qRT-PCR data with qDATA. For Case 2, average Ct values were precomputed starting from the TRs of any target gene and endogene three values sets. This approach generates only three ΔCt values for the included categories.

As a first example, we evaluated the gene expression of *LManVI* gene in wild-type Oregon fruit fly males and in mutant *γCOP*^*14a*^/*γCOP*^*14a*^ males [6] using *RpL32* as an endogene. The raw Ct data were used to assess the relative expression of *LManVI* in four experimental male groups, namely not-infected Oregon (OregonNI), not-infected *γCOP*^*14a*^/*γCOP*^*14a*^ (14NI), infected Oregon (OregonInf) and infected *γCOP*^*14a*^/*γCOP*^*14a*^ (14Inf). For the infection assay, *P. aeruginosa* inoculum mixed with sucrose solution was delivered by ingestion.

Figure 4 shows the tabular results of the two-way two sample Student’s t-tests performed on 2^-ΔCt^ values by qDATA fed with the input tables corresponding to Case 1 and Case 2 approaches. For Case 1 (Figure 4A), the results reveal statistical significance between each of the sample groups, with t values ranging between -9.435 and +2.451 and p values within the 0.017 -7.44 × 10^−13^ interval. In Case 2 (Figure 4B), the statistical significance is dimmed, as only one comparison has p < 0.05. These results suggest that working with only three 2^-ΔCt^ values per BR is not always enough to get a similar statistical significance with the Student’s t-test, even if the null hypothesis has actually to be rejected. Moreover, the ANOVA analysis and post-hoc Tukey HSD test revealed that Case 2 still lacks statistical significance for either of the tests.

**Figure 4.**
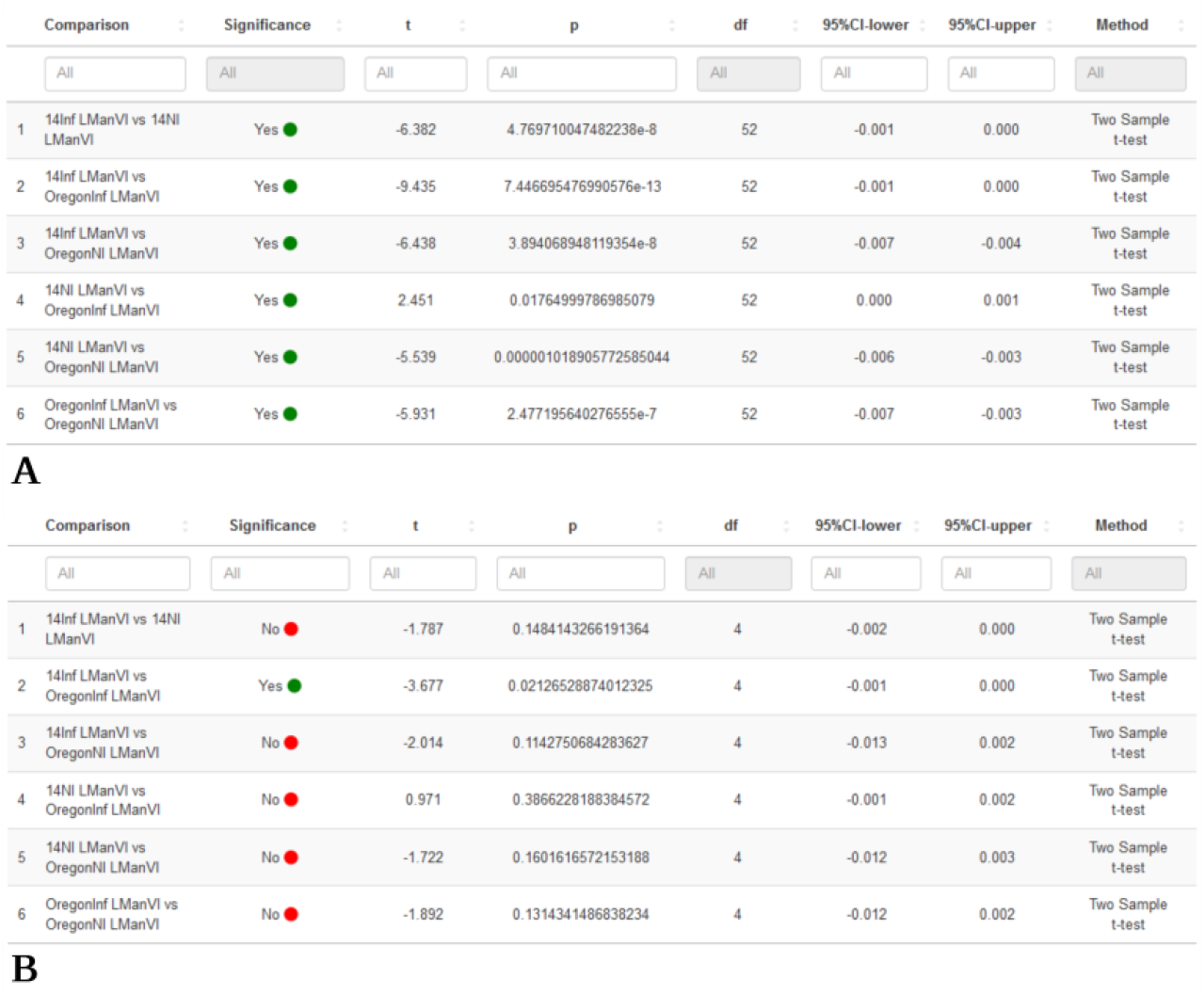
Results of statistical analysis on data corresponding to Case 1 (**A**) and Case 2 (**B**) scenarios. Two-way two sample Student’s t tests were performed on 2^-ΔCt^ values derived from data gathered for each considered fruit fly male groups. Green and red dots highlight the statistical significance. Individual plots were generated in qDATA and figure edits were created with BioRender.com (accessed on 11 November 2023).

FC analysis and log_2_FC revealed notable differences between Case 1 and Case 2 procedures, for both FC values and their corresponding SE values, where the latter one is calculated as mean FC ± SD/√n (n stands for the number of tested individual 2^-ΔΔCt^ values). It has value to mention that even small changes in mean FC could translate into significant variations of the GOI relative expression. In our study, we found that the 14Inf versus OregonInf generates an FC value of 0.524 for Case 1, while the same comparison generates an FC value of 0.188 for Case 2. This result shows that the calculations concerning the relative gene expression may be seriously affected by the input data fed to a statistical test. Here, the individual FC values dataset is 729 for Case 1 and 9 for Case 2. As previously argued, a larger dataset generates a more accurate estimate of the true FC between groups, and it also positively impacts the variations in the SE values, as the SE in Case 1 < SE in Case 2. A similar trend was observed for the log_2_FC values, which represent the log linear form of FC values.

In the second example, we started from data acquired for gene expression of *Relish* and *Toll* relative to the HK gene *Actin* in either summer or winter *A. mellifera* individuals [8]. The qRT-PCR experiments were performed separately, with each gene of interest and the HK gene being analyzed in different 96-well plates. For this reason, we present the corresponding data as two distinct sets, one for *Relish* and another for *Toll*, symbolized qRT1 and qRT2, respectively. Similarly to the previous example, we used two scenarios for feeding data to qDATA, namely Case 1, which handle the complete raw Ct values, and Case 2 that uses the mean Ct values for each TRs category.

Two sample two-way Student’s t tests were performed between each group of interest and the results are shown in Figure 5. As in the fruit fly example, the Case 1 comparisons provide highly statistically significant results, with p values ranging between 0.01 and 6.9 × 10^−14^. Case 2 comparisons led to a weaker significance of the comparisons results, most probable because of a much smaller dataset (df = 52 for Case 1 and df = 4 for Case 2).

**Figure 5.**
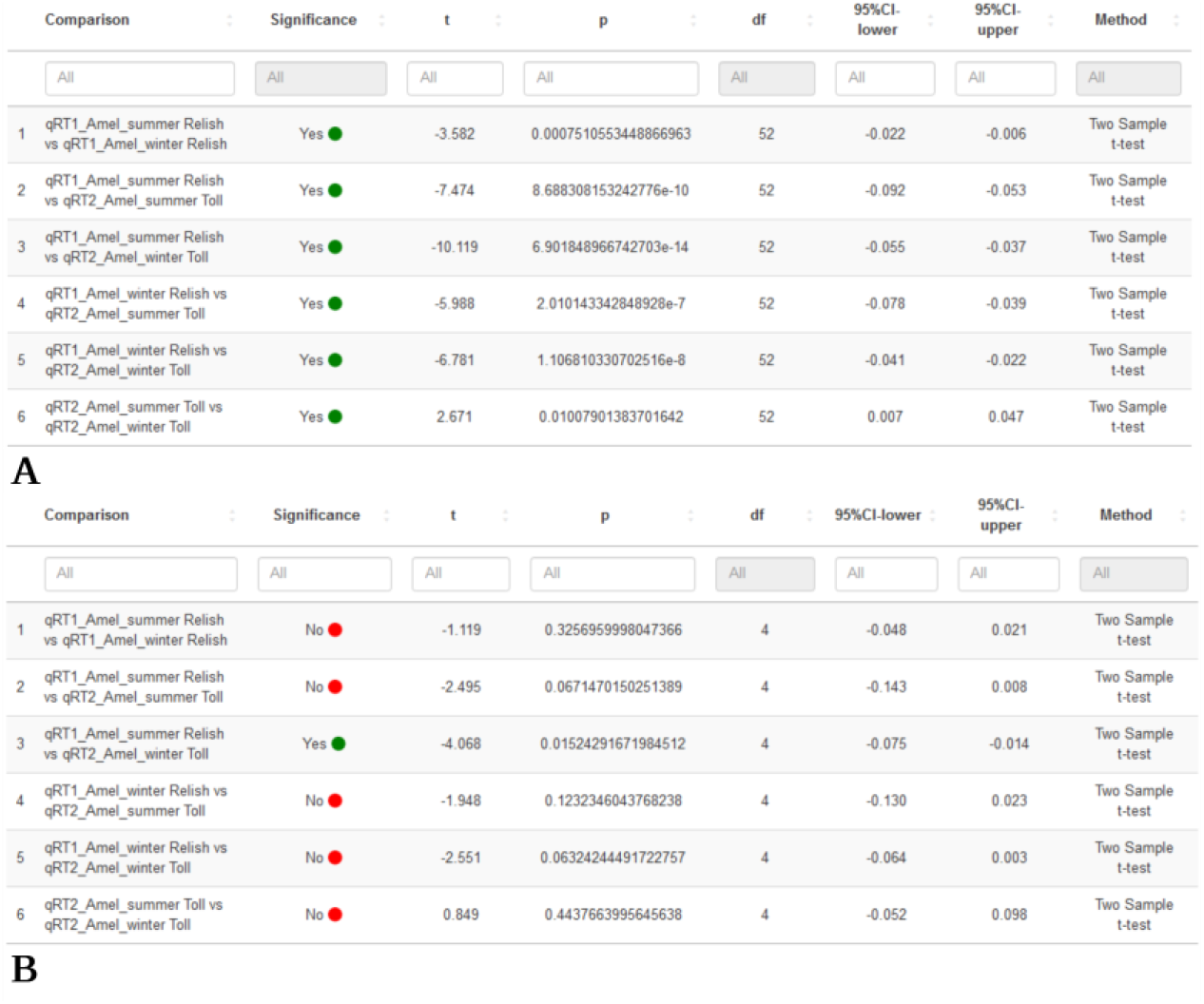
Results of statistical analysis on data corresponding to Case 1 (**A**) and Case 2 (**B**) scenarios. Two-way two sample Student’s t tests were performed on 2^-ΔCt^ values derived from data of each considered *A. mellifera* target gene. Green and red dots highlight the statistical significance. Individual plots were generated with qDATA and figure edits were created with BioRender.com (accessed on 11 November 2023).

Intriguingly, one-way ANOVA results were statistically significant for both case studies. Case 1 dataset generated an F value of 36.474 and a p value of 3.45 × 10^−16^, whereas Case 2 dataset generated an F value of 4.211 and a marginally significant p value of 0.046.

For the *A. mellifera* data, the FC computations did not show large variations between Case 1 and Case 2 datasets, as opposed to the fruit fly data. When considering the same type of comparison, the consistency of results is demonstrated by finding relative gene expression levels with differences of less than 10% between Case 1 and Case 2 approaches. Nonetheless, this aspect shows that even if in specific situations the mean FC estimates are similar regardless of the available individual FC values, their statistical significance is altered when smaller sets of 2^-ΔCt^ values are used.

As demonstrated by the two previous examples, our procedure can accommodate data generated through various experimental setups. The first example involves results concerning the immune response of a eukaryotic host to bacterial challenges. Such studies are becoming very popular since the antimicrobial resistance (AMR) emerged as a major threat to public health worldwide, affecting both patients and socio-economic aspects [9]. The AMR natural phenomenon is artificially accelerated by the selective pressure exerted by irresponsible and exaggerated antibiotic use [10], thus the need for constantly testing the resistance and antibiotic susceptibility using *in vitro* or *in vivo* models. The use of experimental model organisms such as *D. melanogaster* is particularly relevant for *in vivo* infection and antibiotic treatment due to the possibility of performing gene expression analysis concerning both the host and the pathogen of interest, but also the commensal microbiota. Accordingly, increasingly complex experimental designs aim to accomplish these testing. By being able to interpret a flexible input table, qDATA allows scaling qRT-PCR experiments for accommodating a large number of GOI genes, a necessary feature when gene expression data are sought after in microbiology or immunology experimental setups.

A number of software implementations have been developed to automatize various qRT-PCR calculations, as described by Pabinger et al. (2014). Such tools tackle Ct calculation, normalization or quantification, statistics based on Ct values, melting curve analysis, digital PCR result analysis [11]. Most of these tools exist as standalone R packages, Windows or Web dependent, but there are also implementations based on Perl, MATLAB, Java, Python or SAS [11]. Currently, most tools are either no longer maintained or outdated.

We compared qDATA with two recently developed tools, namely the Python based Auto-qPCR [12] and the R package qPCRtools [13]. Auto-qPCR is available either online at https://auto-q-pcr.com/ and requires an active internet connection or offline, requiring python or command prompt skills. qPCRtools is available offline as an R package and can be used only by users skilled in R programming. Conversely, the GUI implementation within the shiny package of qDATA does not require any additional effort from the user. Both Auto-qPCR and qPCRtools require complex input data formats that must contain information about all 96 wells of a qRT-PCR plate and this issue is poorly addressed in their respective software manual. Additionally, Auto-qPCR takes the PCR generated file as input, which might cause issues with some particular qRT-PCR machines. Our implementation does not restrain the user to import all of the data generated in a qRT-PCR experiment, as outliers can be freely removed or multiple experimental plates can be included in the input file. Considering the output graphs, both Auto-qPCR and qPCRtools create basic graphs with instances of overlapping labels, while qDATA generates ready for publication boxplots, violin plots, histograms, and bar plots. Additionally, qDATA builds tables of ΔCt, 2^-ΔCt^ or 2^-ΔΔCt^ values that can be directly imported into other bioinformatics and statistics tools, such as GraphPad Prism, IBM SPSS, Stata, MATLAB and others.

Moreover, neither of the discussed tools considers a clear delimitation between BR and TR, which is a different approach compared with our strategy. qDATA allows the user to customize the data according to the experimental approaches. For example, the user may consider either multiple BRs with multiple TRs or directly the means of TRs, but also only a BR with multiple TRs and so on. Our software aims to be flexible and to accommodate a variety of experiments, which translates into almost no special requirements for the input files. qDATA is a user-friendly implementation of the Livak method that can be run without prior knowledge in bioinformatics or programming. As a perspective, we intend to implement various add-ons to qDATA, such as the accommodation of the Pfaffl mathematical modelling [14], computation of primer efficiency, melting curve analysis or data analysis corresponding to other RT-PCR methodologies.

## 4. Materials and Methods

### 4.2. Software developing

qDATA was developed in the *Drosophila* Laboratory, Faculty of Biology, University of Bucharest, between March and November 2023. The application was written in the R programming language [15] using the RStudio IDE [16]. The application makes use of R core functions and of various installable packages, namely ggplot2 [17], dplyr [18], tidyr [19], shiny [20], shinythemes [21], DT [22], gridExtra [23], ggthemes [24], nortest [25], tseries [26], reporter [27], qqconf [28], qqplotr [29], shinycssloaders [30], shinybusy [31], shinycustomloader [32], car [33], rstatix [34], fostawesome [35], RColorBrewer [36], ggtext [37], coin [38] and exactRankTests [39].

Implementations for running qDATA on Linux OS were performed in bash on Linux Mint 21.1 Vera MATE Edition and on Ubuntu 22 LTS Jammy Jellyfish. Both OSs were run in virtual environments with Oracle VM Virtual Box.

### 4.2. Data collection

The data used in the two described examples were generated in the *Drosophila* Laboratory. Data analysis and results interpretation were previously published [6,8]. Input data used for the analysis presented in the article is not provided here.

### 4.3. Cross-validation

The results generated by qDATA were cross-validated by performing manual or automatic computations in Microsoft Excel or GraphPad Prism version 8.0.2 for Windows (GraphPad Software, Boston, Massachusetts USA, www.graphpad.com), respectively.

## 5 Patents

qDATA is freely available on GitHub where it has received copyright under the Creative Commons Zero v1.0 Universal (CC0-1.0 Universal) license.

## Supplementary Materials

Not yet applicable.

## Author Contributions

Conceptualization, A.C.R., A.A.E. and A.I.; methodology, A.C.R., A.A.E. and A.I.; software, A.I.; validation, A.C.R. and A.I.; formal analysis, A.C.R. and A.I.; investigation, A.C.R., A.A.E. and A.I.; resources, A.C.R., A.A.E and A.I.; data curation, A.C.R., A.A.E., A.I., M.C.C.; writing—original draft preparation, A.C.R. and A.I.; writing—review and editing, A.C.R., A.A.E., A.I. and M.C.C.; visualization, A.C.R. and A.I.; supervision, A.C.R.; project administration, A.C.R., M.C.C.; funding acquisition, M.C.C. All authors have read and agreed to the published version of the manuscript.

## Institutional Review Board Statement

Not applicable.

## Informed Consent Statement

Not applicable.

## Data Availability Statement

qDATA can be downloaded at GitHub at https://github.com/A-Ionascu/qDATA.

## Acknowledgments

We thank Nicoleta Denisa Constantin for the relevant feed-back on the article’s text and the help and support on the subject of software implementation in Linux OS.

